# Lipid nanodiscs facilitate the identification of a fragment compound inhibiting the enzymatic activity of the bacterial membrane protein MsbA

**DOI:** 10.1101/2021.06.15.448612

**Authors:** Kaoru Fujimoto, Akinobu Senoo, Satoru Nagatoishi, Daisuke Takahashi, Tadashi Ueda, Kouhei Tsumoto, Jose M.M. Caaveiro

## Abstract

Membrane proteins are critical elements of numerous therapeutic approaches ranging from cancer to bacterial infections. MsbA is a bacterial membrane protein that has received increasing attention as an antibacterial target for its role in the processing of Lipid A, a key precursor of lipopolysaccharide that is essential for bacterial growth. When employing nanodiscs it is possible to stabilize MsbA by providing a membrane-like environment that enhances its enzymatic activity. Taking advantage of this property we have carried out a fragment screening using the biophysical method of surface plasmon resonance. This approach identified several compounds that bind specifically to MsbA. In particular, one of these fragment molecules not only binds to the target, but also inhibits the ATPase activity of the MsbA protein. The similarity of this fragment to the adenine moiety of ATP points at a route to generate stronger and more potent inhibitors for MsbA and even other proteins of its family of ABC transporters. Collectively, our study reveals biophysical approaches that facilitate the identification of fragment candidates inhibiting the activity of membrane proteins.

## INTRODUCCION

Membrane proteins are responsible for critical cellular functions such as signal transduction, energy production, and metabolite transport. Because of their importance for cellular function and homeostasis, membrane proteins such as G protein-coupled receptors or ion channels are often drug targets for therapeutic intervention [1]. In this respect, the emergence of bacterial resistance is a topic of growing concern. The development of new antibiotics is urgently required, still, in recent years antibiotics with new mechanisms of action have been scarce [2]. Novel targets, including novel membrane protein targets, and more aggressive approaches to drug discovery are sought after to address these serious concerns.

MsbA is a dimeric ABC transporter expressed on the inner membrane of the Gramnegative bacterium *Escherichia coli* (*E. coli*) that transports lipid A, a necessary precursor of lipopolysaccharide (LPS, also known as endotoxin) [3–5). Translocation of Lipid A from the inner leaflet to the outer leaflet of the inner membrane by MsbA is powered by the enzymatic hydrolysis of ATP at the cytoplasmic side and coupled to large-scale conformational changes. Because Lipid A is essential for bacterial growth, MsbA has recently attracted attention as a target molecule for new antibacterial agents [6, 7]. In addition, since ABC transporters play a major role in bacterial drug resistance and pharmacokinetics in humans, the discovery of inhibitors/modulators of MsbA, and the understanding of their inhibition mechanisms may facilitate the elucidation of the activity and regulatory mechanisms of other ABC transporters [8].

The introduction of lipid nanodiscs for the manipulation and stabilization of membrane proteins have greatly benefited their study [9], and MsbA is not an exception [10]. Such nanodiscs keep membrane proteins in a native-like state, since nanodiscs mimick many properties of natural membranes [11], and thus avoiding some of the shortcoming when employing detergents. The usefulness of nanodiscs is demonstrated in a recent study in which a large library of three million compounds was employed to identify novel inhibitors of MsbA by classical high-throughput approaches [6].

Although routinely employed by large pharmaceutical companies, high-throughput approaches are very challenging for academic research units, which favor a more focused efforts such as that of fragment-based screening [12]. Fragment compounds, which are the building blocks of small-molecule drugs, have the advantage of small molecular weight (less than 300 molecular weight) covering a large chemical space with small libraries of a few hundred or thousand compounds [13]. Because small molecular weight (generally) means weaker binding energy to the target protein, the detection of favorable interactions relies on highly sensitive biophysical methodologies, particularly X-ray crystallography and nuclear magnetic resonance (NMR) [14]. These methods require large amounts of protein sample, and long periods for screening, restricting the applicability of fragment methods to membrane proteins. The development of a more expeditious screening method that can respond to the characteristics of membrane proteins (minute samples) is highly sought after.

The technique of surface plasmon resonance (SPR) offers a good compromise between speed and demand of protein resources that, in several aspects, potentially rival or even surpass X-ray crystallography or NMR [15]. However, the use of SPR for fragment screening, and in particular that of membrane proteins, has been limited until recently because of the intrinsic difficult to detect the minute binding signals of fragment compounds, and the limited choices to stabilize membrane proteins for the entire length of the experiments. Novel experimental approaches and equipment is relieving that situation in a progressive fashion [16, 17].

Although high-throughput screening of MsbA in nanodiscs using enzymatic assays has been used to identify compounds for inhibitors of MsbA [6], there have been no similar reports employing biophysical techniques. In this study we aimed to develop search novel inhibitors of MsbA using fragment-based approaches by SPR. We combined the synergistic effect of nanodiscs as a model reproducing the lipid bilayer environment, and SPR which provides a good balance between sensitivity and the costs in terms of protein resources. We reveal several small fragments binding to MsbA, one of them inhibiting the ATP activity of MsbA. The nature of the inhibitor is such that may suggest a plausible strategy for the optimization of its potency.

## MATERIALS AND METHODS

### Cloning, expression and purification of MsbA

MsbA was amplified by PCR from genomic DNA of *E. coli* K12 strain with forward primer CGTG<u>GGATCC</u>CATAACGACAAAGATCTCTCTACGTGGCAG and reverse primer CAGC<u>AAGCTT</u>TCATTGGCCAAACTGCATTTTGTGAAGTTGC containing the BamH I and Hind III sites (underlined), and cloned into a pET26 vector with a His6 tag at the C-terminus. The protein was expressed in *E. coli* C43 (DE3) transformed with the MsbA expression vector prepared above (Kan+). Protein expression was induced with 1 mM isopropyl-β-D(−)-thiogalactopyranoside. Four hours after induction, the bacteria were harvested by centrifugation at 4,000 × *g* for 10 min at 4 °C. Cell pellets were resuspended in 40 mM Tris, 300 mM NaCl, 2 mM dithiothreitol (DTT) at pH 8.0, followed by cells lysis with a French press at 1 kBar. The supernatant recovered after centrifugation (25,000 × *g*, 1 hour, 4 °C) was subjected to a round of ultracentrifugation at 150,000 × *g* for 1 hour at 4 °C in buffer supplemented with fresh DTT at 1 mM. The precipitated fraction was resuspended in 40 mM Tris, 300 mM NaCl, 1 mM DTT, 5 mM imidazole, and solubilized in the presence of 1% (w:w) n-dodecyl-β-*D*-maltoside (DDM) for 2 hours at 4 °C. The resulting supernatant was collected by ultracentrifugation (150,000 × *g*, 1 hour, 4 °C) and applied onto an immobilized metal-ion affinity chromatography column (Talon, Clontech) equilibrated with 40 mM Tris, 300 mM NaCl, 1 mM DTT and 0.1% (w:w) DDM. MsbA was eluted with the same buffer supplemented with 300 mM imidazole. The eluted fractions containing MsbA were collected and concentrated to 100-200 μM by ultrafiltration (Amicon Ultra, Millipore), and the samples frozen in liquid nitrogen, and stored at −80 °C.

### Expression and purification of MSP1

Expression vector of membrane scaffold protein 1 (MSP1) displaying an N-terminal His7 and TEV cleavage site were expressed and purified as described previously [10, 18]. TEV protease containing a His6 tag at the N-terminal was purified using a Ni^2+^-affinity column as described elsewhere [19]. MSP was treated with TEC protease, and the cleaved MSP fragment separated from non-cleaved MSP and TEV protease in a HisTrap HP column (Cytiva). The fractions of the flow-through containing MSP protein were pooled together and concentrated with a 10 kDa Amicon Ultra-filter (Millipore) to 100-200 μM, and stored at stored at −80 °C until use.

### Determination of lipid concentration

To determine the concentration of lipid we employed the method of Barlett [20]. Samples of phospholipid DMPC were digested with 400 μl of 72% (v:v) perchloric acid for 2 hours at 180 °C. Samples were diluted with 4.2 ml of water, and treated with 200 μl of 5% (w:w) ammonium molybdate and 200 μl of freshly prepared amidol solution (0.1 g amidol, 2 g sodium metabisulfite, 10 ml H_2_O). The mixture was incubated in a boiling water bath for 10 minutes. Since one molecule of DMPC contains one atom of phosphorous. the concentration of DMPC was determined by comparison with a standard curve prepared with phosphoric acid solution.

### Preparation of lipid

The appropriate amount of lipid 1,2-dimyristoyl-sn-glycero-3-phosphocholine (DMPC) in chloroform was aliquoted into test tubes and dried under reduced pressure in a draft to completely remove the chloroform. The dried lipid as dissolved in 50 mM HEPES, 200 mM NaCl, pH 7.4, and 100 mM sodium cholate to a lipid concentration of 50 mM, by stirring and sonication.

### Preparation of nanodiscs

A mixture of lipid DMPC at 50 mM in nanodisc buffer (50 mM HEPES, 200 mM NaCl, pH 7.4) supplemented with 14-40 mM, scaffold protein MSP1, and MsbA dissolved in nanodisc buffer was stirred gently at 4 °C for 1 hour. Subsequently the sample was incubated with BioBeads SM-2 at 0.6 g/ml to remove excess surfactant by gently rotating at 23 °C for 4-6 h. BioBeads were then removed by centrifugation. Nanodiscs containing MsbA were further purified on a Superdex 200 increase 10/300 GL column (GE healthcare) equilibrated with nanodisc buffer.

### Measurement of ATPase activity of MsbA

The enzymatic activity of MsbA (ATP hydrolysis) was based on the method of Doerrler et al. [3]. MsbA (5 μg) solubilized in 0.1% DDM or embedded in nanodiscs was incubated with 250 μl of enzyme buffer (100 mM HEPES, 20 mM MgCl2 at pH 7.4) and various concentrations of ATP (and othr molecules when necessary) were adjusted to a volume of 500 μl and incubated at 37 °C for 10 min. For MsbA solubilized in DDM, the assay buffer was supplemented with 0.2 % DDM. To the reaction solution, 150 μl of 12 % (w:w) SDS was added to stop the reaction. To quantify the phosphorous produced during the hydrolysis of ATP into ADP and phosphate, 300 μl of a solution containing 6 % (w:w) ascorbic acid and 1 % (w:w) ammonium molybdate in 1 M HCl prepared within the last 24 h was added and incubated at room temperature for 3-6 min. To this 450 μl of a solution containing 2% (w:w) citric acid monohydrate, 2% (w:w) sodium metaarsenite, and 2% (v:v) acetic acid was incubated with the enzyme reaction products for more than 20 min. The absorbance at 850 nm was measured within the next four hours. The amount of phosphoric acid released due to hydrolysis was quantified by comparing the absorbance with that of a standard phosphoric acid solution reacted by the same method, and the ATPase activity of MsbA (nmol/min/μg of MsbA) determined.

### Isothermal titration calorimetry (ITC)

The interaction between nanodisc-embedded MsbA and the ATP analog AMP-PNP was determined in a calorimeter MicroCal Auto-iTC200 (Cytiva). MsbA embedded in nanodiscs and the non-hydrolyzable analog AMP-PNP were and prepared at 20-50 μM and 100-500 μM, respectively. The temperature of the experiment was adjusted to 25 °C. MsbA was loaded into the calorimeter cell and the solution of AMP-PNP was loaded into the titration syringe for measurement. The buffer was 50 mM HEPES, 200 mM NaCl, 5 mM MgCl2 at pH 7.4. Samples of protein were dialyzed against the same buffer. Data were integrated using NITPIC [21, 22] and analyzed with SEDPHAT [23]. Origin 7 (OriginLab, MA) was also used for data analysis.

### Surface plasmon resonance (AMP-PNP)

The binding analysis of the substrate analog AMP-PNP was carried out in a Biacore 8K instrument (GE Healthcare) using an NTA Series S sensor chip (Cytiva). MsbA reconstituted in nanodiscs was immobilized to the flow cell by the His-tag at the C-terminal, whereas empty nanodiscs (no MsbA present) was immobilized to the reference cell by the His-tag at the N-terminus of the protein MSP1. The His-tag of MSP1 was cleaved with TEV protease prior to the preparation of nanodiscs containing MsbA. The binding experiment was carried out in 50 mM Tris, 100 mM NaCl, 10 mM MgCl2, 0.005% Tween 20, and 5 % DMSO with increasing concentrations of ligand from 0.31 to 20 μM in two-fold serial dilutions at 25 °C. The ligand was flowed into the sensor chip at a rate of 30 μ/min for 60 second and a dissociation time of 300 seconds. Bulk solvent correction was applied automatically using reference solutions containing 4.5, 5.0, 5.5. and 6.0% DMSO. The signal in the flow cell (MsbA-nanodiscs) was subtracted with the signal in the reference cell containing empty nanodisc. The difference between the flow and reference cells was taken as the indication of specific binding to MsbA. Data analysis was carried out with the Insight Evaluation software provided by the manufacturer of the instrument (GE Healthcare).

### Surface plasmon resonance (screening)

Fragment screening of MsbA embedded in nanodiscs was performed with the Discovery-ZEN-LIBRARY 1 (Zenobia) in a Biacore 8K instrument (GE Healthcare) using an NTA Series S sensor chip (Cytiva) at 25 °C. Nanodiscs with and without MsbA were immobilized as described above. In each run, fragments were flowed at 30 μ/min for 30 in in 50 mM Tris, 100 mM NaCl, 10 mM MgCl2, and 0.005% Tween 20 and 5% DMSO. Bulk solvent correction was performed as above. Because the immobilization of MsbA to the NTA chip is not covalent, a small amount of MsbA nanodiscs is dissociated from the surface as the experiment progressed, and therefore the screening was divided in two halves. In between them a fresh sensor chip with MsbA-nanodiscs immobilized was necessary. The signal in the flow cell (MsbA-nanodiscs) was subtracted with the signal in the reference cell containing empty nanodisc. We note that prior to the screening with MsbA, a clean screen was performed in which the fragments were evaluated with empty nanodiscs. A total of 17 compounds were found to strongly interact with the empty nanodiscs and were not deployed in the screen with MsbA-embedded nanodiscs. Data analysis was carried out with the Insight Evaluation software provided by the manufacturer of the instrument (GE Healthcare). Fragments in the top 10% responses were selected for concentration-dependent changes in the signal intensity. Fragments were flowed at increasing concentrations from 31 to 250 μM in two-fold dilution series. The difference was calculated as the signal of the compound binding to MsbA.

## RESULTS and DISCUSSION

### Preparation of MsbA in nanodiscs

Previous studies have demonstrated the usefulness of nanodiscs as a platform for the analysis of membrane proteins in general [9], and MsbA in particular [6, 10]. Herein we prepared nanodiscs using MSP1 and DMPC based on previous studies. In order to increase the yield of nanodisc formation in our experimental set-up, we optimized the concentration of cholate to be near its critical micellar concentration (14 mM at pH 7.5) at the time of mixing MsbA with MSP1 and lipid. The purification of nanodiscs by SEC shown in Figure S1 indicated that the yield approached 90%, a satisfactory value for the overall study. In the same figure, and consistent with the same reports, it is observed that the enzymatic activity (ATP hydrolysis) of MsbA is clearly promoted in nanodiscs with respect to that of MsbA in DDM. These preliminary data clearly support the use of nanodiscs for the characterization of MsbA (see below).

### Biophysical characterization of the interaction between MsbA and a substrate analog

In order to prepare the experimental set up for the screening of an inhibitor (see below), we employed a non-hydrolyzable compound (AMP-PNP) and examine its inhibitory effect, as well as its binding affinity by two biophysical methodologies (Figure 1). The inhibitory AMP-PNP effect of AMP-PNP was evaluated at a concentration of 250 μM. The Michaelis-Menten plot shows a decrease in the ATPase reaction rate. The analysis of the kinetic parameters *K_M_* and *V_max_* indicated that the value of *K_M_* increased three-fold, consistent with a mode of competitive inhibition, a mechanism that could have been anticipated from the significant similarities with the actual substrate ATP.

**Figure 1:**
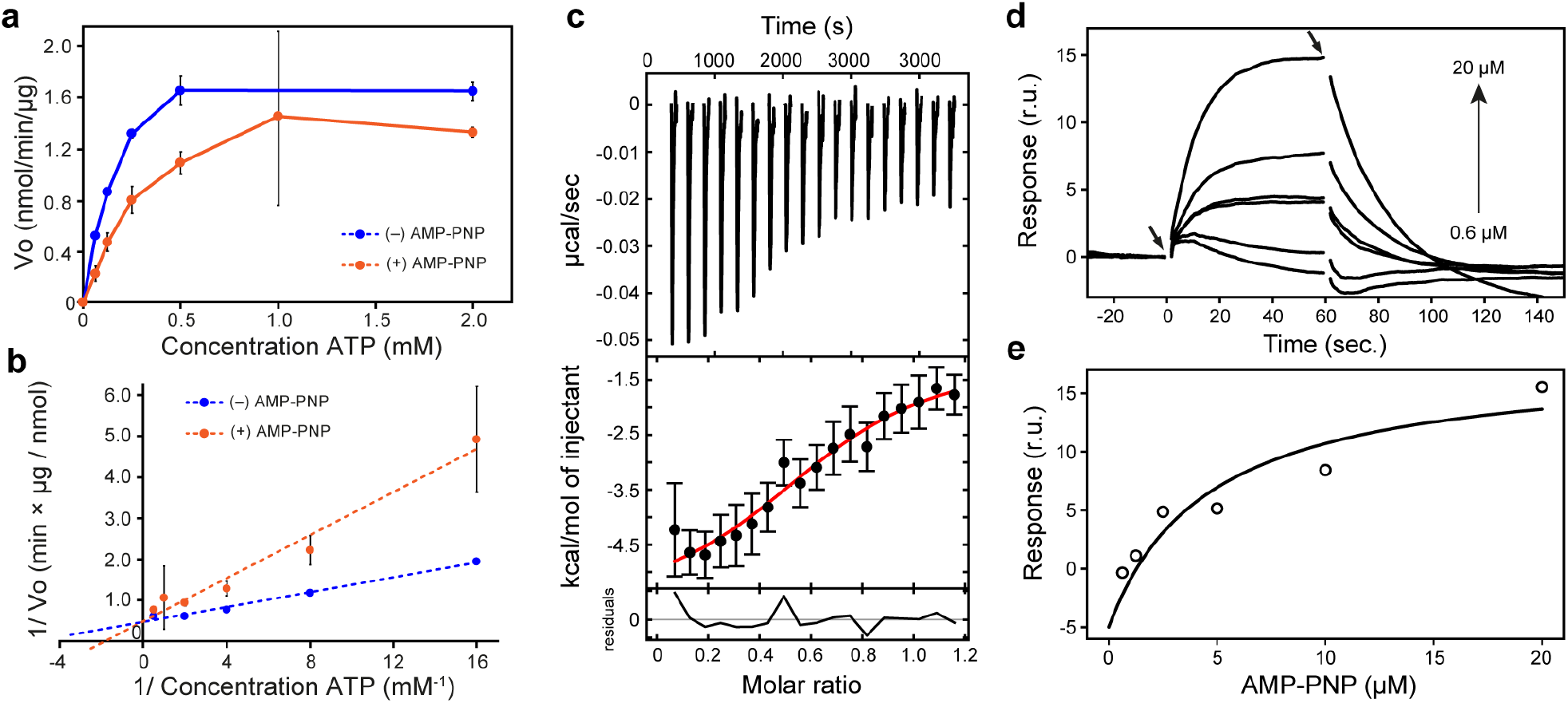
Biophysical characterization of the binding of substrate analog AMP-PNP to MsbA in nanodiscs. **(a)** Enzymatic activity of MsbA in the absence (blue) or presence (orange) of 250 μM AMP-PNP. The data corresponds to the average of three determinations ± SE. Often the error was contained within the symbol. **(b)** Double reciprocal representation to determine the values of the kinetic parameters *K_M_* (0.21 ± 0.18 mM without compound; 0.61 ± 0.28 nM with compound) and *V_max_* (2.21 ± 0.18 nmol min^−1^ μg^−1^ without compound; 2.06 ± 0.74 nmol min^−1^ μg^−1^ in the presence of AMP-PNP). **(c)** Binding of AMP-PNP examined by ITC. The top and bottom panels represent the titration curve and the binding isotherm, respectively. The analysis yielded the equilibrium constant (*K_D_* = 5.1 μM), change of enthalpy (ΔH = −4.9 kcal mol^−1^) and change of entropy (ΔS = 7.7. cal mol^−1^ K^−1^). Analysis was carried out with the programs NITPIC and SEDPHAT as described in material and methods section. **(d)** Binding sensorgrams by SPR. Serial two-fold dilutions of AMP-PNP were injected to a surface decorated with MsbA embedded in nanodiscs. **(e)** Binding isotherm obtained by computing the binding response at the end of the period in which the compound was injected (before 60 sec.). A binding constant of 4.6 μM was determined.

To determine the binding affinity of AMP-PNP to MsbA in nanodiscs, something not reported before, two experiments were envisioned. First, we carried out a determination of the binding constant by the calorimetric method of ITC, the golden standard in interaction analysis [13]. The titration of MsbA with AMP-PNP gave ride to small exothermic peaks (Figure 1c). The analysis of the data with the programs NITPIC and SEDPHAT revealed that the equilibrium constant was in the μM range (*K_D_* = 5.1 μM) driven by a modest but favorable changes of enthalpy (*ΔH* = −4.9 kcal mol^−1^) and entropy (−*TΔS* = −2.3 kcal mol^−1^).

Next a critical experiment was performed for this study, consisting in the evaluation of the binding of the same AMP-PNP to the same MsbA-embedded nanodiscs by SPR. The comparison with the ITC data was critical to assess the merit of the technique for a screening with a library of compounds. In the sensorgrams, the binding of the substrate analog to the membrane protein gave rise to an increase of response units in a dose-dependent fashion (Figure 1d,e). Although a clear downwards drift of the signal was present in the response cause by nanodisc leak, it was satisfactorily corrected during the analysis phase by the software provided by the manufacturer. In some cases, for example the curve obtained at a concentration of 20 μM analog, some abnormal trace is observed during the second half of the dissociation step. In the sensorgram corresponding to the lowest concentration tested (0.3 μM) a small but negative signal was recorded, which may be caused among other things by a swing in the concentration of DMSO. The difference in DMSO concentration is corrected by solvent correction module to minimize bulk effects. When the correction range is exceeded, the reliability of the response obtained by the anolyte binding decreases. In addition, MsbA is immobilized in the active cell whereas empty nanodiscs were immobilized in the reference cell, and although their response signals are subtracted, there were times when the response appears to be negative due to differences in the decrease in the immobilized amount in each cell.

These difficulties did not influence the determination of the equilibrium constant, since the calculation was not based on the kinetic phases, but in the maximal response at the end of the association phase. A binding constant of 4.6 μM was determined, a value remarkably similar to that determined by ITC, thus validating our SPR experimental approach.

### Screening of MsbA embedded in nanodiscs with a fragment library by SPR

In this study, we aim at screening a library of fragments of small molecular weight (average molecular weight 200 Da) against a nanodisc-embedded membrane protein. The total molecular weight of the MsbA/lipid/MSP1 ensemble is estimated at 220 kDa, *i.e*. about 500 times greater that the size of the fragments (considering MsbA is a dimer), explaining in part why fragment screening by SPR methods using nanodisc-embedded membrane proteins has been difficult so far. Since there is a lack of accumulated empirical data for fragment methodologies using SPR under these combined conditions, we were very interested in the results of this research.

Prior to the actual screen, a control experiment was carried out. In this preliminary assay, all 352 fragments contained in the library were sequentially injected to evaluate their suitability for the actual screen. A total of 15 compounds were eliminated from the list of candidates, including those that greatly reduced the amount of immobilized nanodiscs and those that strongly bound to empty nanodiscs before and after injection of the anolytes.

For this primary screen, MsbA nanodiscs were immobilized in the active cell and empty nanodiscs in the reference cell. The difference in the signals between flow and reference cell for each of the 337 compounds (distributed in two groups as explained in Materials and Methods) is plotted in Figure 2a. When analyzed case-by-case, the types of responses could be classified into three groups Figure 2b. A large group of fragments yielded little or no increase of signal, whereas in other cases a moderate increase of signal within the limit of response expected for a small molecule. Finally, a third category of fragments producing large responses were observed in a few cases. Because due to the general low affinity of the fragments, it was possible that they bind to multiple locations, so even large responses could not be discarded without further evidence, and therefore we decided to select the top 10% (33 compounds) with the highest signal intensity as hit candidates in the primary screening.

**Figure 2.**
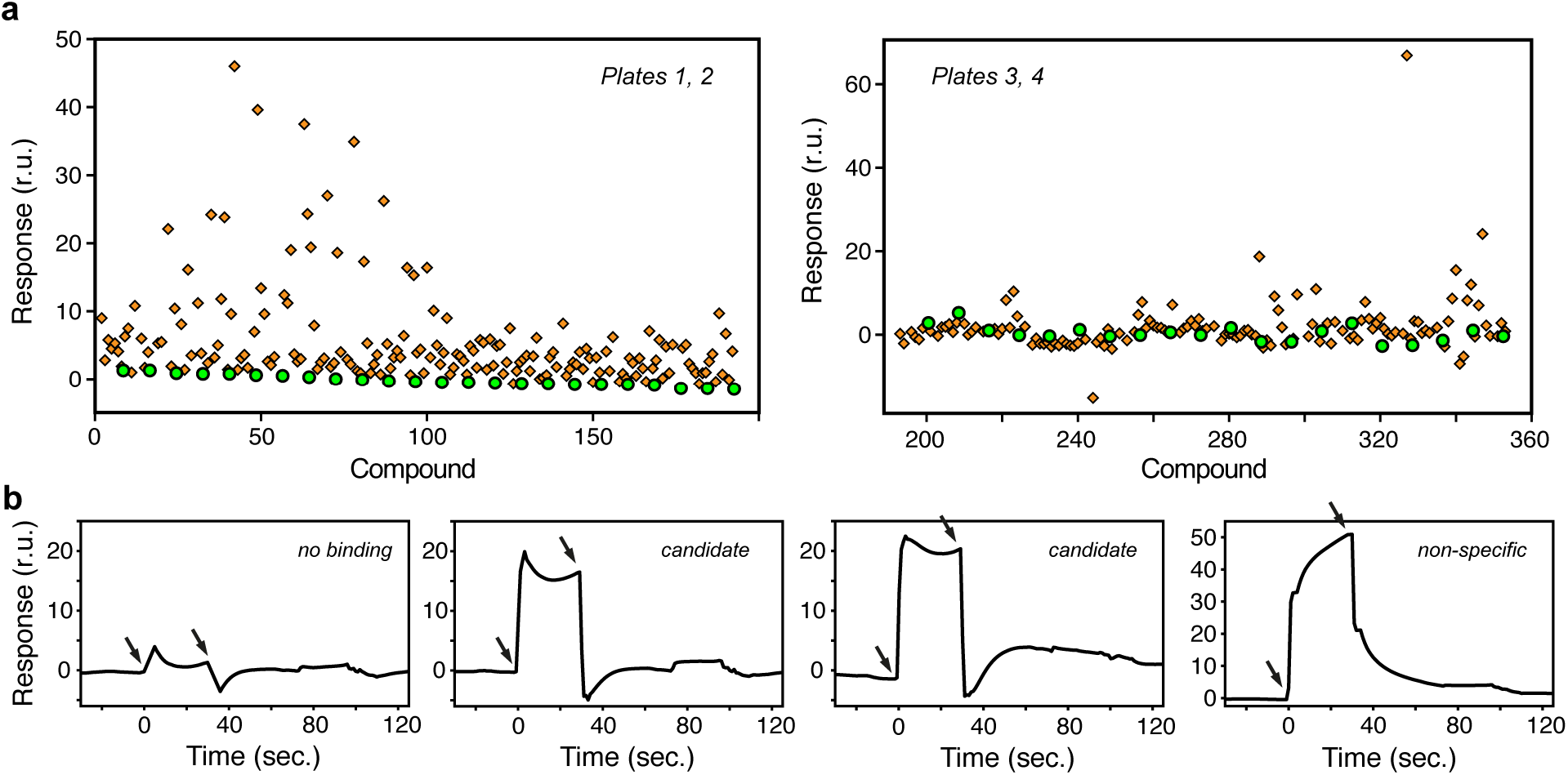
Screening of MsbA embedded in nanodiscs with a fragment library by SPR. **(a)** MsbA was screened with a fragment library comprising 360 compounds (Discovery-Zen-Library 1) by SPR. Data of plates 1 and 2, and of plates 3 and 4 are shown on the left and right panels, respectively. Each compound was allowed to contact MsbA immobilized in a His-NTA chip for 30 seconds. The orange diamonds correspond to the signal obtained in the presence of fragment compounds, whereas the green spheres to control experiments. **(b)** Representative SPR sensorgrams that illustrate some of the typical responses in the experiment. The first arrow indicates injection of the compound, and the second arrow injection of buffer. Fragments that yielded sensorgrams with little change of signal (left panel) were not considered further. Middle panels exemplify two putative candidates, and the right panel an example of a likely unspecific binding (large response value).

### Dose dependence and validation

The 33 compounds selected from the primary screening were cross-validated by examining their concentration-dependent responses by SPR (Figure 3a and Figure S2a-b). The binding responses of three fragments (12, 24 and 57, their structures are included in their respective figures) increased with the fragment concentration and were compatible with specific binding, i.e. producing moderate response values. In contrast, most of the fragments either displayed responses not commensurate with the increasing concentration, or their responses were too large compared with the expected response values for a molecule of their size (Figure S2c). The three fragments indicated above (12, 24 and 57) were selected as the final candidate compounds. Although the calculation of the dissociation constant was not reliable (high error) for fragments 24 and 57, the data suggested it to be in the sub-milimolar range. The *K_D_* value for compound 12 was 78 μM.

**Figure 3.**
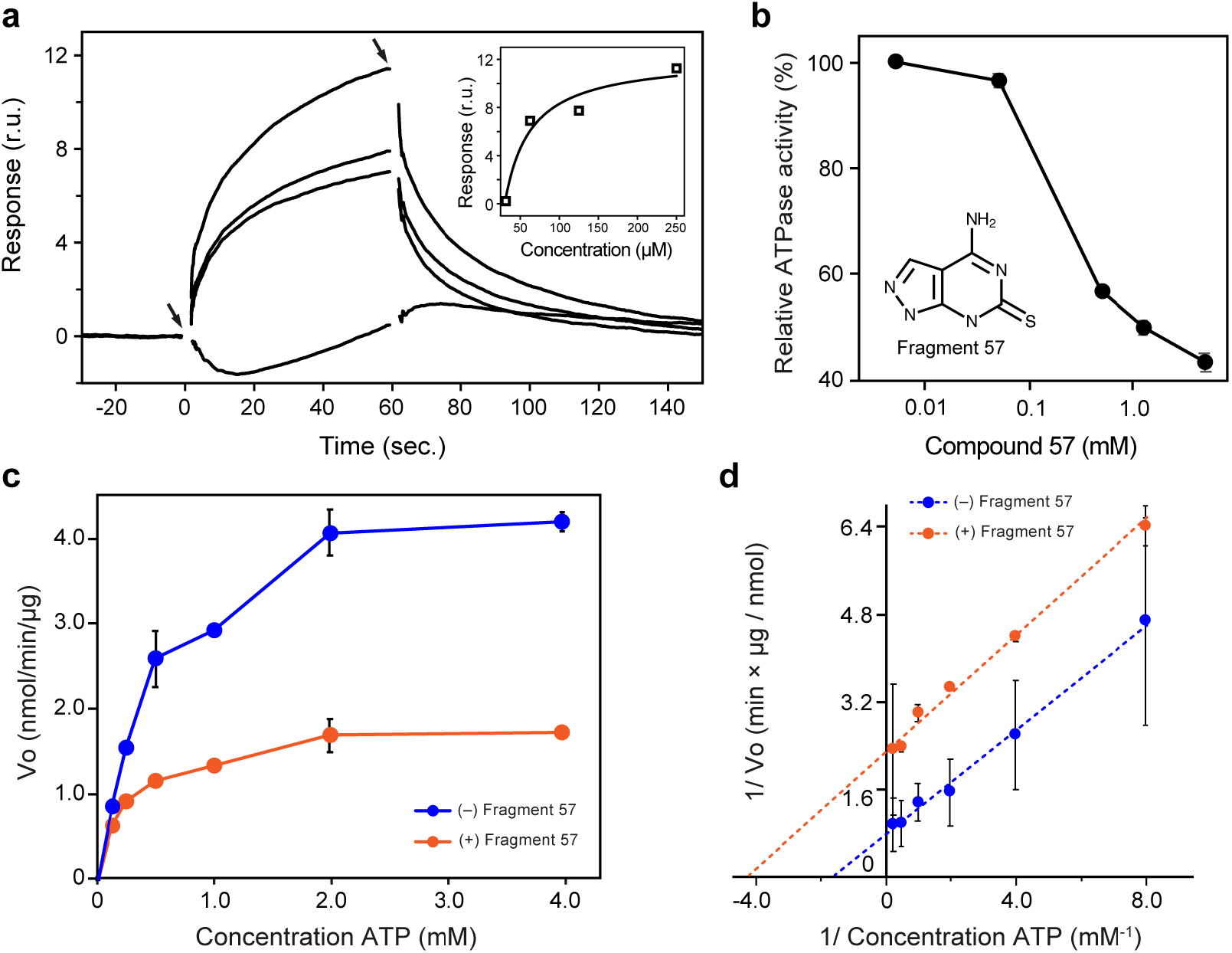
Characterization of a hit candidate (fragment 57). **(a)** SPR response of nanodiscs of MsbA in the presence of increasing concentrations of fragment 57. The binding response was compatible with specific binding of this compound to the membrane protein. The inset shows the binding isotherm from which as *K_D_* in the sub mM was determined. Since the uncertainty of this determination was significant, no value is given here. **(b)** Inhibition of the ATPase activity of MsbA in the presence of increasing concentrations of fragment 57. The figure shows the relative activity of the enzyme in the presence of fragment with respect to the absence of it. The structure of compound 57 is shown in the inset. **(c)** Comparison of the enzymatic activity of MsbA in the presence (orange) or absence (blue) of fragment 57. **(d)** Double reciprocal plot to determine the kinetic parameters *K_M_* (0.62 ± 0.06 mM without compound; 0.35 ± 0.19 nM in the presence of fragment 57) and *V_max_* (5.12 ± 0.34 nmol min^−1^ μg^−1^ without compound; 1.60 ± 0.34 nmol min^−1^ μg^−1^ in the presence of fragment 57). For panels (b~d) the data corresponds to the average of three determinations ± SE. Often the error bar was contained within the symbol.

One of the drawbacks of a screening of these characteristics is that the binding of the fragments to the protein does not ensure a biological effect, or an inhibitory role. In general, the activity of the ATP hydrolysis domain of the ABC transporter is inhibited, resulting in blocked substrate transport into and out of the cell. Thus, we evaluated their biological inhibitory potential by the same ATPase enzymatic assay employed above for AMP-PNP (Figure S2d). Only fragment 57 showed a significant degree of inhibition of the enzymatic activity of MsbA, whereas fragments 12 and 24 showed little or essentially no deleterious effect for the ATPase activity even at the high concentration selected for this assay (up to 7.5 mM). On the contrary, fragment 57 showed a characteristic dose-dependence behavior at concentrations between 5 μM and 5 mM (Figure 3b).

The inhibitory effect was further quantified by enzymatic analysis. A three-fold decrease in the reaction rate (*V_max_*) catalyzed by MsbA in the presence of fragment 57 with respect to the protein without fragment was observed (Figure 3c,d). The structure of fragment 57 is similar to that of adenine, a component of ATP (Figure S3), suggesting that fragment 57 binds directly to the same pocket in the ATP binding site. However, the low value of *V_max_* value suggests that Fragment 57 exerts its inhibitory activity by a mixed mechanism other than pure competitive inhibition.

### Structure activity relationship

In an attempt to increase the potency of the inhibitor, and also to explore if a structural homolog of fragment 24 could inhibit the ATPase activity of MsbA, we obtained nine compounds from commercial sources (Figure S3). To enhance the interaction with MsbA, we selected compounds with various kinds of substituents, such as more donor and acceptor groups for hydrogen bonding, halogen groups, large substituents, and different hydrophobicity. The inhibitory potential of the nine compounds, as well as that of fragment 57, was evaluated at a concentration of 5 mM with 2 mM ATP. None of the new compounds (seven analogs to compound 24, and three compounds similar to fragment 57) exceeded the inhibitory effect of fragment 57. Compound 5 of the group of analogs to fragment 24 showed incipient inhibitory potential in this assay, as well as compound 7.

Another interesting feature resulted from the comparison between the structure of adenine and fragment 57 (Figure 4b). Since adenine is a component of ATP, it was interesting to observe that fragment 57 showed a significantly greater inhibitory potency than an actual portion of the substrate. The effect is clear and suggest that the incorporation of a thioketo group at a specific position enhance the interaction with MsbA. Since ABC transporters use active transport (powered by ATP) this feature may be of general applicability for this large family of proteins and that could be worth considering to prepare and design stronger and more effective inhibitors.

**Figure 4:**
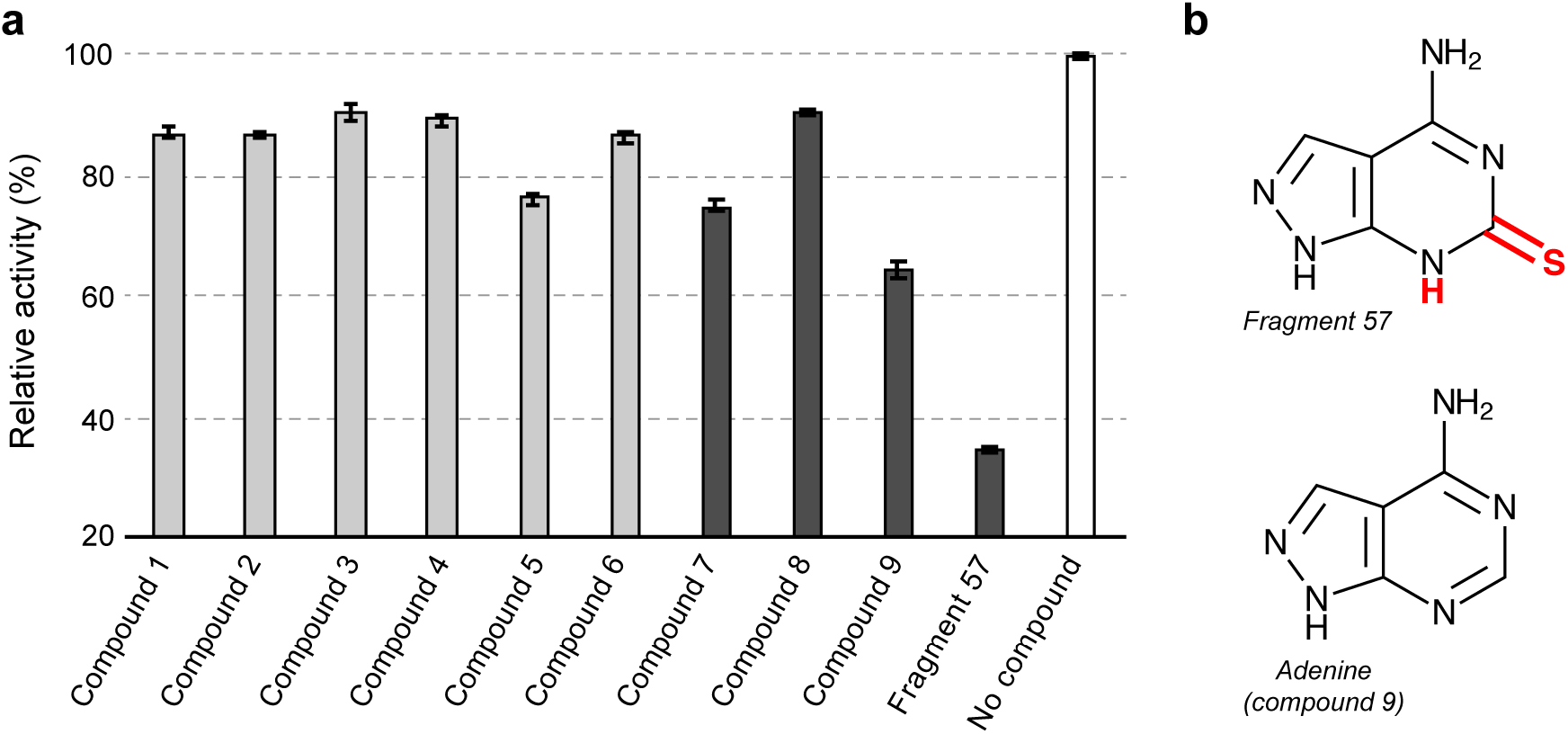
Mini structure-activity relationships experiment. **(a)** Relative enzymatic activity of MsbA in the presence of various compounds with respect to MsbA without compounds. The light and dark gray bars correspond to compounds similar to fragment and fragment 57, respectively. The structure of these compounds is shown in Figure S3. The white bar corresponds to the 100% activity obtained with MsbA in the absence of compounds. The data corresponds to the average of three determinations ± SE. (b) Structures of fragment 57 and compound 9 (adenine). Since adenine is part of the natural substrate ATP, this result suggests that the presence of a thioketone group (red) attached to adenine increases the interaction of the nucleobase with the protein.

## Conclusion

In this study we have employed nanodiscs to stabilize the membrane protein MsbA and evaluated its usefulness to carry a fragment screening employing the broadly employed biophysical methodology of SPR. To the best of our knowledge this is the first example of a fragment screening of a membrane protein embedded in nanodiscs by SPR. We found that the use of a positive control (the substrate analog AMP-PNP) was a fundamental piece for the successful optimization of the conditions of the experiment. The screening and further cross-validation identified three fragment molecules with clear binding propertied to MsbA. One of them (fragment 57) showed clear inhibitory effect, and moreover, from the comparison of its structure to that of adenine we could suggest a route for the design of compounds of higher inhibitory potency.

Collectively, we have established that SPR is a suitable to method for a membrane protein stabilized by nanodiscs, using MsbA as an example. We hope that similar approaches will be applied to other membrane proteins embedded in nanodiscs.

## Abbreviations

*E. coli*: *Escherichia coli*
LPS: Lipopolysaccharide
DTT: Dithiothreitol
DDM: n-dodecyl-β-D-maltoside
DMPC: 1,2-dimyristoyl-sn-glycero-3-phosphocholine
MSP1: Membrane scaffold protein
AMP-PNP: 5’-adenylimiimidodiphosphate
*E. coli*: *Escherichia coli*
SPR: Surface plasmon resonance,
ITC: Isothermal titration calorimetry,
TEV: Tobacco etch virus
SEC: Size exclusion chromatography
SE: Standard error
DMSO: Dimethyl sulfoxide

## Acknowledgements

This work was supported by the Platform Project for Supporting Drug Discovery and Life Science Research, Basis for Supporting Innovative Drug Discovery and Life Science Research [BINDS] from AMED JP21am0101091 (to T.U. and J.M.M.C.) and JP21am0101094 (to K.T.).

