## Supplementary data for "Lipid nanodiscs facilitate the identification of a fragment compound inhibiting the enzymatic activity of the bacterial membrane protein MsbA"

*[1]Laboratory of Protein Structure, Function, and Design, Graduate School of Pharmaceutical Sciences, Kyushu University, 3-1-1 Maidashi, Higashi-ku, Fukuoka 812-8582, Japan; [2]Department of Chemistry and Biotechnology, Graduate School of Engineering, The University of Tokyo, 7-3-1 Hongo, Bunkyo-ku, Tokyo 113-8656, Japan; [3]Institute of Medical Science, The University of Tokyo, 4-6-1 Shirokanedai, Minato-ku, Tokyo 108-8639, Japan; [4]Department of Bioengineering, School of Engineering, The University of Tokyo, 7-3-1 Hongo, Bunkyo-ku, Tokyo 113-8656, Japan; [5]Laboratory of Global Healthcare, Graduate School of Pharmaceutical Sciences, Kyushu University, 3-1-1 Maidashi, Higashi-ku, Fukuoka 812-8582, Japan,*

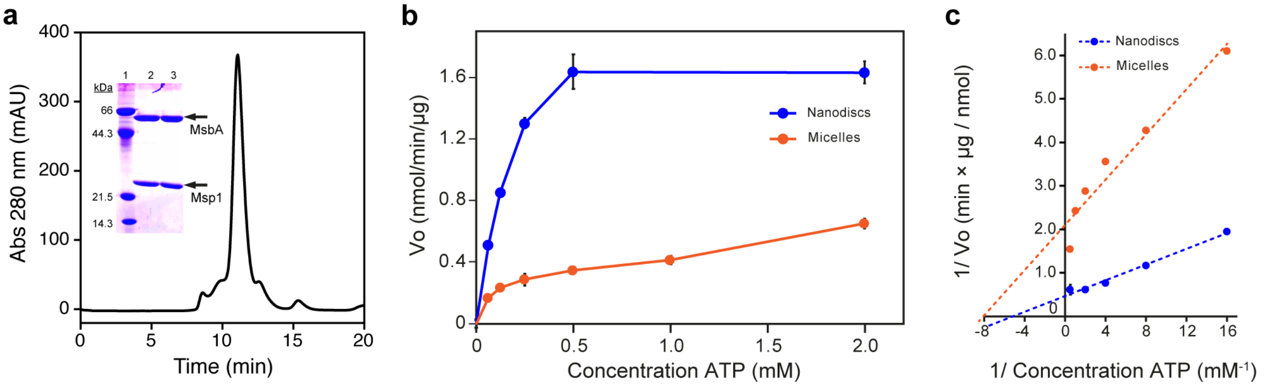


**Figure S1. Lipidic nanodiscs efficiently preserve the enzymatic activity of MsbA. (a)** Nanodiscs containing MsbA were separated from insoluble material by SEC. The large peak contains MsbA embedded in nanodiscs of MSP1 and lipid. The inset corresponds to an SDS-PAGE gel of the main peak (lanes 2-3). The molecular weight markers appear in lane 1. The position of MsbA and Msp1 are indicated by arrows. **(b)** Enzymatic rate (ATPase activity) of MsbA in nanodiscs (blue) is compared with that in micelles of DDM (orange) under the same experimental conditions. The data corresponds to the average of three determinations ± SE. Often the error was contained within the symbol. **(c)** Double reciprocal analysis to determine the values of the kinetic parameters $K_M$ (0.21 ± 0.18 mM in nanodisc; 0.14 ± 0.03 mM in micelles) and $V_{max}$ (2.21 ± 0.18 nmol min$^{-1}$ µg$^{-1}$ in nanodiscs; 0.47 ± 0.06 nmol min$^{-1}$ µg$^{-1}$ in micelles).

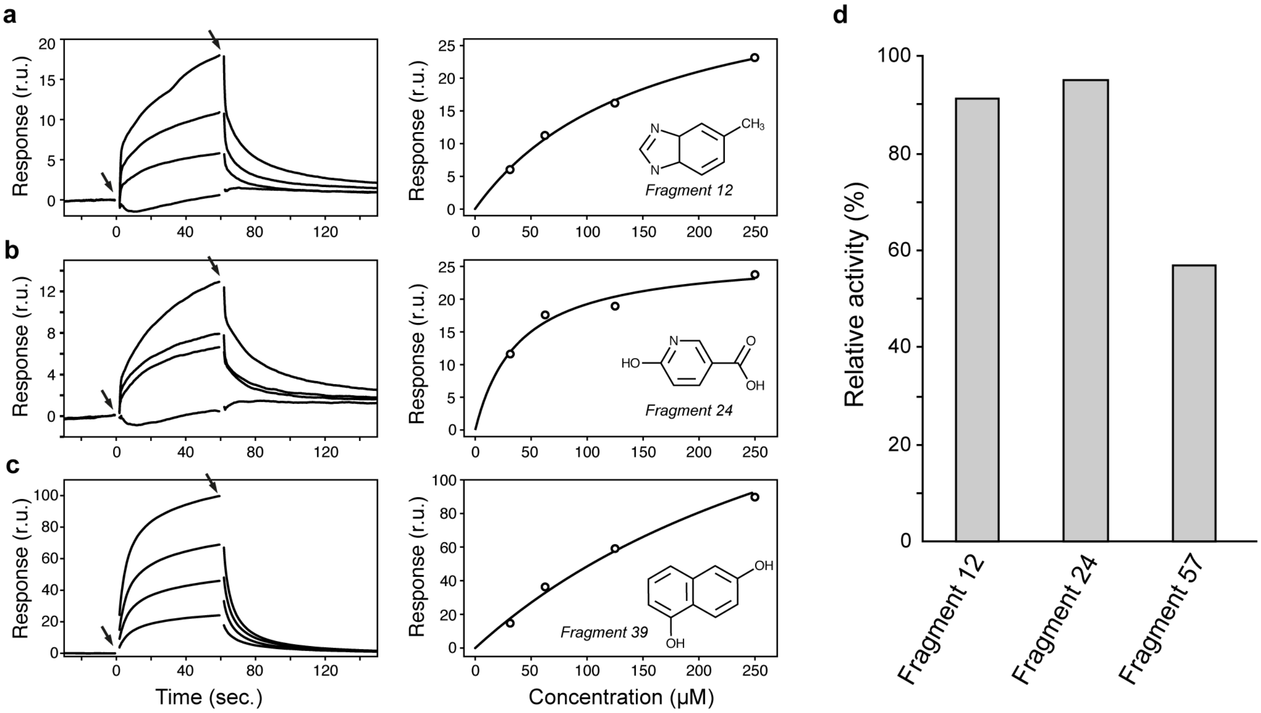


**Figure S2. Binding of fragments to MsbA.** Sensorgrams (left panels) and binding isotherms (right panels) of nanodiscs containing MsbA in the presence of fragment 12 **(a)**, fragment 24 **(b)**, or fragment 39 **(c)**. Whereas fragments 12 and 24 showed signs of specific binding, fragment 39 clearly exceeded the maximal response signal for a fragment of its size (non-specific binding) and was not further considered as a binding candidate. The structures of the fragments are given inside the right panels. To facilitate the comparison of the three binding isotherms, the signal was shifted to start at zero. This cosmetic effect does not have any influence in the calculation of the binding constants, which was done separately. **(b)** Relative enzymatic activity of MsbA in the presence of the indicated fragments (at 5 mM) with respect to the activity of protein in their absence using 2 mM ATP. Only fragment 57 showed a significant degree of inhibition to MsbA among the three candidates tested.

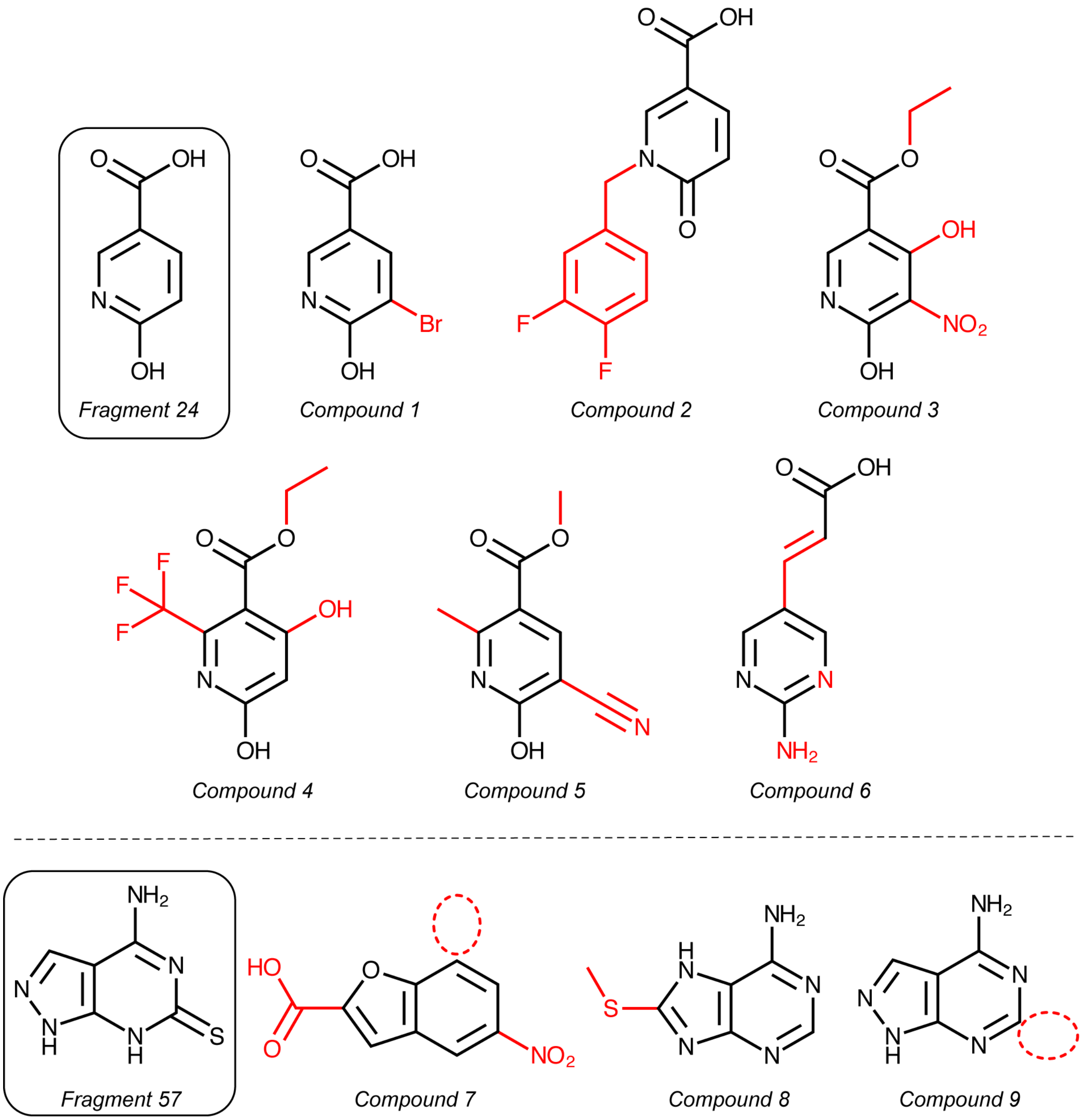


**Figure S3. Families of compounds employed in the structure-activity relationship assays.** Compounds above and below the dashed line are related to fragment 24 and fragment 57, respectively.
